# Neutralizing Antibodies with Neurotropic Factor Treatment Maintain Neurodevelopmental Gene Expression Upon Exposure to Human Cytomegalovirus

**DOI:** 10.1101/2023.03.02.530870

**Authors:** Benjamin S. O’Brien, Rebekah L. Mokry, Megan L. Schumacher, Suzette Rosas, Scott S. Terhune, Allison D. Ebert

## Abstract

Human cytomegalovirus (HCMV) is a beta herpesvirus that causes severe congenital birth defects including microcephaly, vision loss, and hearing loss. Infection of cerebral organoids with HCMV causes significant downregulation of genes involved in critical neurodevelopmental pathways. The precise features of the infection causing this dysregulation remain unknown. Entry of HCMV into human cells is determined by the composition of glycoproteins in viral particles, which is influenced by the source of the virus. This includes a trimer complex and a pentamer complex with the latter enriched from replication in epithelial cells. To begin dissecting which features contribute to neuronal pathogenesis, we evaluated infection using virus from different sources along with the distribution of cellular entry receptors on cells in cerebral organoids. We observed significant increases in the number of viral genomes, viral spread and penetrance, and multinucleated syncytia in neural tissues infected with HCMV propagated in epithelial cells compared to fibroblasts. To determine if this was related to entry receptor distribution, we measured expressions of cellular entry receptors and observed similar distributions of all receptors on cells obtained from organoids indicating that source of virus is likely the key determinant. Next, we asked whether we could limit pathogenesis using neutralization antibodies. We found that pre-treatment with antibodies against viral glycoprotein B (gB) and gH successfully decreased viral genome levels, viral gene expression, and virus-induced syncytia. In contrast, targeting specific cellular entry receptors failed to limit infection. Using an antibody against gB, we also observed partial protection of developmental gene expression that was further improved by the addition of brain derived neurotropic factor (BDNF). These studies indicate that source of HCMV is a key determinant of neuronal pathogenesis that can be limited by neutralization antibodies and neurotropic factors.

## INTRODUCTION

Human cytomegalovirus infection is the leading cause of non-heritable birth defects worldwide and remains a serious threat of morbidity for immunocompromised individuals. Although inhibitors of viral DNA replication are available, adverse side effects limit their use in pregnant people (1) thereby limiting adequate treatment options for this highly vulnerable patient population. A congenital HCMV infection occurs when the virus is passed to the developing fetus during pregnancy, which happens in roughly 1:200 live births in the US (2, 3). The most serious cases cause lifelong neurological abnormalities such as microcephaly, sensorineural hearing loss, and developmental motor delays (3-5). In the United States 40-80% of individuals experience infection by age 40 (6, 7). Fortunately, symptoms are typically mild for a healthy individual, and the virus will eventually enter a latent stage with occasional reactivations remaining in the body (3, 6). This life-long persistence has been associated with diverse diseases such as immuno-senescence, atherosclerosis, gliomas, and Alzheimer’s Disease.

One reason for HCMV’s high prevalence is because of its permissive nature infecting many different cell types including endothelial cells, epithelial cells, fibroblast, lymphocytes, macrophages, and neural cells (8, 9). Until recently, however, most studies characterizing viral infection were conducted using non-neuronal model systems leading to questions regarding entry into neural cells and the mechanisms by which the virus disrupts normal neurodevelopmental pathways. What is clear is that HCMV infection in neural progenitor cells and cerebral organoids causes massive transcriptional downregulation of several developmentally critical transcription factors, dysregulation of calcium signaling and action potential generation, and disruption of cortical layer development and neural rosette formation (10-12). However, it remains unclear what specific viral genes or proteins cause the neurodevelopmental abnormalities or whether viral entry is a necessary part of the mechanism. Our previous work showed that full viral replication was not required to induce transcriptional and functional abnormalities in neural tissues and that downregulation of key developmental genes was not fully dependent on immediate-early (IE) 1/2 expression (11). Therefore, here we focused on better understanding what role viral entry plays in neurodevelopmental changes.

Previous studies have outlined a mechanism showing that viral entry relies on interactions between viral glycoproteins and host cell receptors expressed at the plasma membrane. As with other herpes viruses, HCMV entry requires viral glycoproteins gB and gH/gL in which gH/gL acts as the host receptor binding protein and gB has a role in fusion and pH-dependent entry (13, 14). Specifically, the gH/gL glycoprotein can form a disulfide-linked trimer with glycoprotein O (gO) to form the trimeric complex (TC) (15-17) or it can unite with viral proteins UL128, 130, and 131 to form the pentameric complex (PC) (13). Additionally, a recent unbiased screening of potential HCMV viral receptors identified a list of host receptors that likely interact with the TC (NRG2, PDGFRα, TGFβRIII) or PC (CD46, Nrp2, FCAR, LILRB3) (18). TC interactions are reported to be required for entry into fibroblasts, whereas epithelial entry requires TC and PC interaction (9, 13).

In these studies, we focus our attention on the influence of HCMV entry in neuropathogenesis. We evaluate the contribution of different sources of virus known to influence virion glycoprotein composition on HCMV entry and replication in neuronal tissues along with entry receptor distributions. Further, we test the effects of neutralizing antibodies and the neurotrophin factor BDNF on protecting neurodevelopmental gene expression upon HCMV challenge.

## RESULTS

### HCMV propagated on epithelial cells exhibits increased tropism for cerebral organoids

Human iPSC-derived cerebral organoids were generated from a healthy control iPSC line (19, 20) and infected after 30 days (30d) of development using a BAC-derived recombinant HCMV strain TB40/E expressing eGFP. Viral stocks were generated on either on epithelial cells, referred to as TB40-BAC4_epi_ or fibroblasts, referred to as TB40-BAC4_fib_. We infected organoids using 500 infectious units per µg of tissue (IU/µg) due to the natural variance in size and cell number in developing organoids. During infection, organoids were placed on a rocker for 2 hr, inoculum was then removed, washed with PBS, and then allowed to progress for 14 dpi. We observed GFP fluorescence by 4 dpi, and this signal continued to increase over the subsequent 10 days indicating viral spread through the tissue (**Fig. 1A,B**). Florescence was substantially higher in organoids infected with the epithelial-derived virus, TB40-BAC4_epi_ compared to TB40-BAC4_fib_. To quantify these differences, we measured viral genomes relative to a cellular gene observing approximate 3-fold more genomes upon infect by TB40-BAC4_epi_ by 3 dpi (**Fig. 1C**). We also determined the mean fluorescent intensity of projection z-stacks at 14 dpi revealing 10-fold higher florescence when infected using epithelial-derived virus (**Fig. 1D**). These data demonstrate that virus originating from epithelial cells exhibits increased tropism for neuronal tissues and is consistent with past studies involving PC-mediated viral entry.

**Figure 1.**
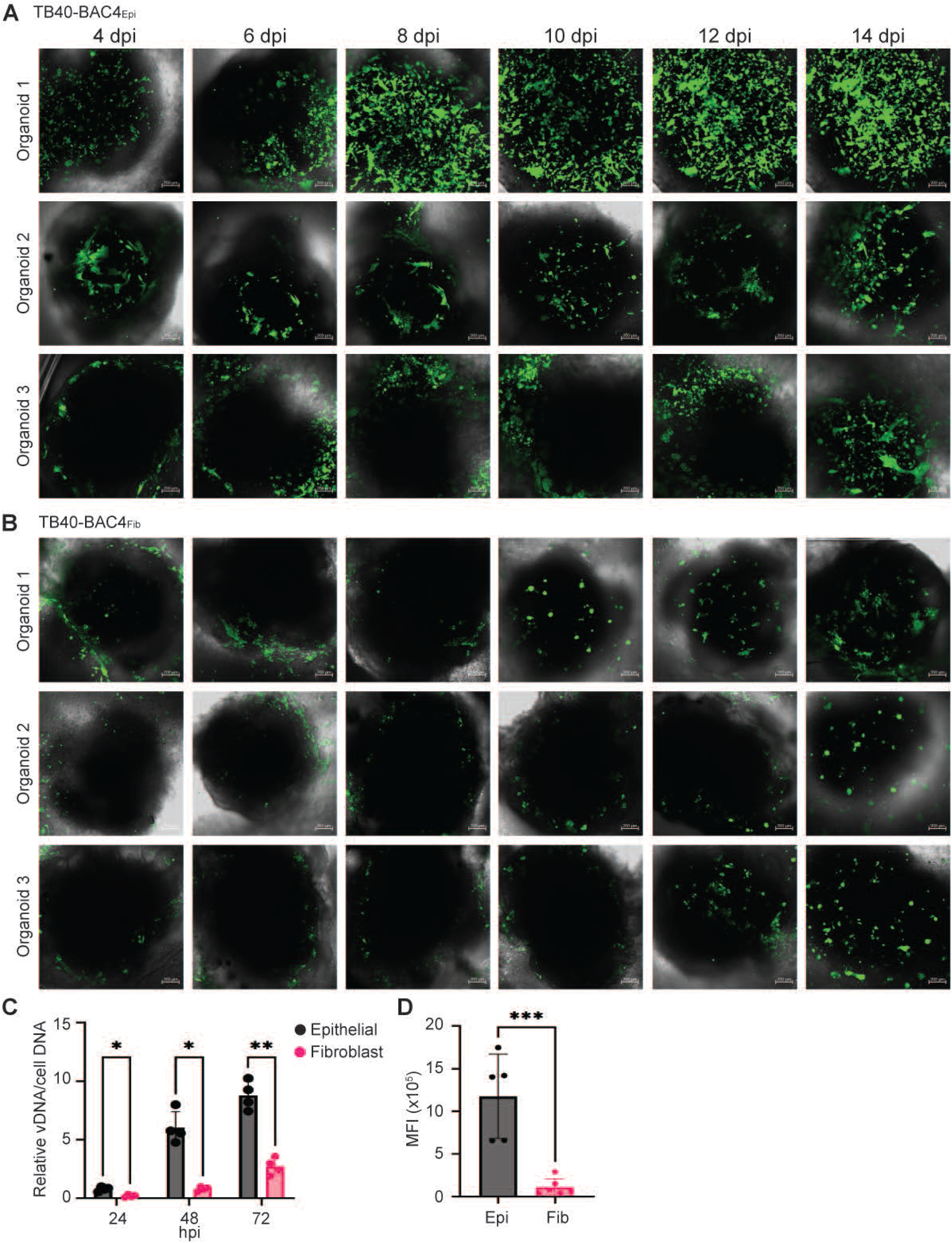
HCMV propagated on epithelial cells exhibits increased tropism for cerebral organoids. Infection of iPSC-derived cerebral organoids at 30 days (30d) of differentiation with HCMV at an MOI of 1 IU/µg using virus produced from MRC5 fibroblasts (TB40-BAC4_Fib_) or ARPE19 epithelial cells (TB40-BAC4_Epi_). **(A)** Overlayed brightfield and GFP images of 3 representative (30d) organoids infected with TB40-BAC4_Epi_ at 4, 6, 8-, 10-, 12-, and 14-days post-infection (dpi) show considerable spread and propagation of GFP signal. Size bar 200 µm **(B)** Overlayed brightfield and GFP images of 3 representative 30d organoids infected with TB40-BAC4_Fib_ at the same time points show the spread and propagation of GFP signal **(C)** Average intensity of GFP fluorescence within Z-stack images of TB40-BAC4_Fib_ or TB40-BAC4_Epi_ infected organoids at 14 dpi. All images were captured at 5x using a Z-stack imaging protocol on a Zeiss LSM9800 microscope. **(D)** Viral genomes measured by qPCR using primers to the UL123 gene relative to GAPDH and whole organoid DNA infected with TB40-BAC4_Fib_ or TB40-BAC4_Epi_. (n=3). Significance was determined using Students T test.

To quantify infection in 3-dimensional space, we measured viral spread and penetrance differences between viral stocks. We rendered the 2D z-stack images taken at 14 dpi into 3D using Zeiss Zen Blue image processing software present representatives for each condition (**Fig. 2A,B**). We have included maximum intensity projection images with heat maps indicating depth of signal penetrance. Infection using TB40-BAC4_epi_ reached an average depth of 650 µm from the organoid surfaces of four organoids compared to TB40-BAC4_fib_ at an average depth of 525 µm (**Fig. 2C**). Penetrance was further quantified by measuring the GFP signal intensity at the centermost z-plane from stacks imaged at 14 dpi which shows significantly higher GFP signal at center slices of TB40-BAC4_epi_ infected tissues (**Fig. 2D**). Finally, we analyzed differences in the 3D space by binning GFP signal intensities into 25 equal bins. We find that organoids infected by TB40-BAC4_epi_ have a higher frequency of voxels within each GFP intensity bin versus TB40-BAC4_fib_-infected organoids (**Fig. 2E**). These studies demonstrate that epithelial-derived virus exhibits greater tropism for neural tissues impacting viral spread and tissue penetrance.

**Figure 2.**
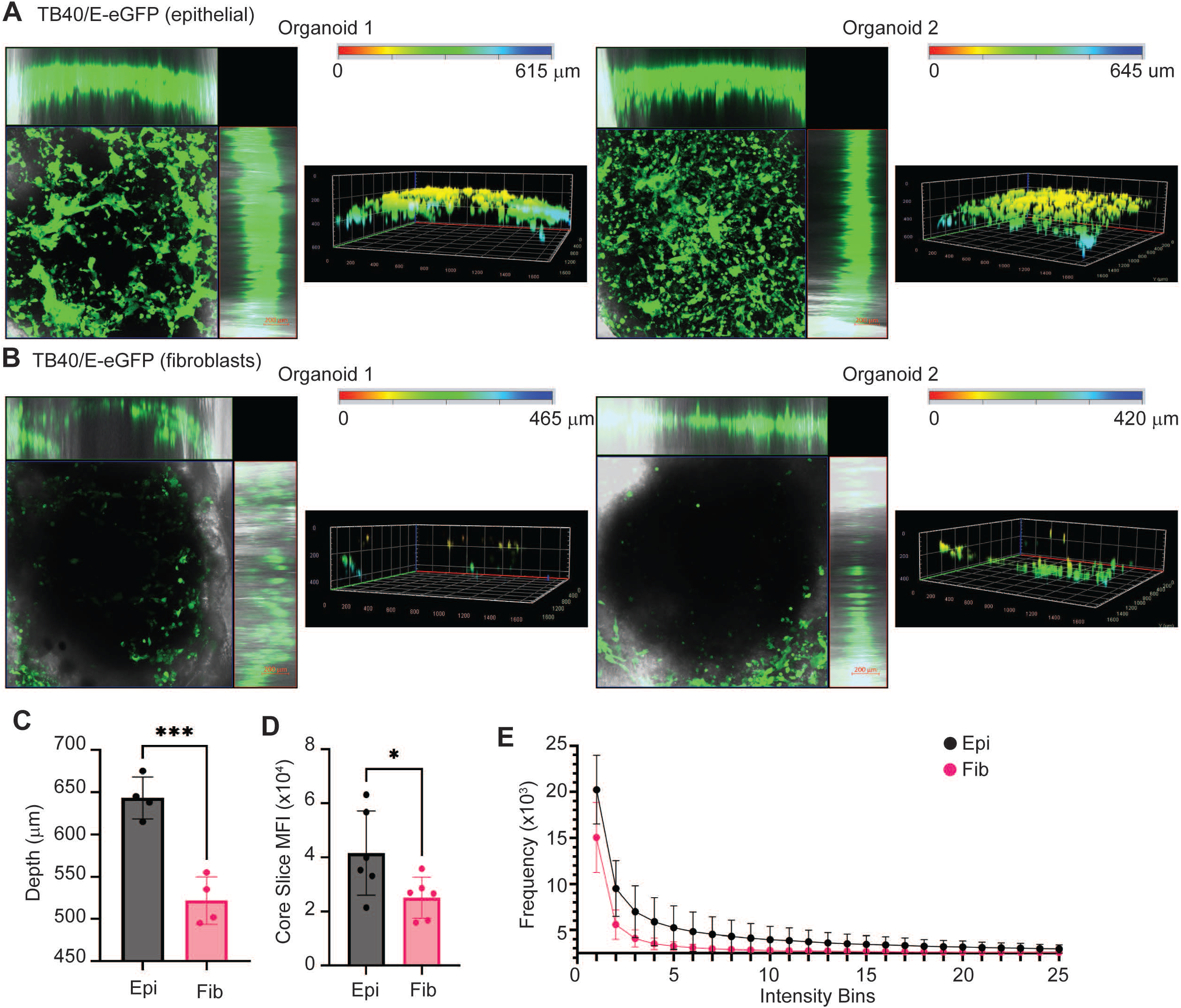
Infection by HCMV TB40-BAC4_Epi_ results greater spread and depth of penetrance compared to TB40-BAC4_Fib_. **(A)** Max intensity projection images along with heat maps of GFP penetrance from two representative TB40-BAC4_Epi_ infected 30d organoids at 1 IU/µg show wide-spread across the surface and penetrance of up to 645 µm. **(B)** Max intensity projection images along with heat maps of GFP penetrance from two representative TB40-BAC4_Fib_ infected organoids with penetrance of up to 465 µm. **(C)** Quantification of deepest depth of penetrance based on rendered images in (A) and (B). All images were captured at 20x using a Z-stack imaging protocol on a Zeiss LSM9800 microscope. **(D)** A measure of the GFP signal at the most central z-plane of TB40-BAC4_Epi_ versus TB40-BAC4_Fib_ infected organoids. **(E)** Binning of the GFP average intensity projection image based on signal intensity (ascending from 1-25) shows TB40-BAC4_Epi_ has an increase in the frequency of pixels falling into the highest intensity bins compared to the TB40-BAC4_Fib_. Five organoid images were analyzed per group at 14 dpi using organoids from 2 differentiations. Significance was determined using Students T test.

### NPCs and cerebral organoids express receptors for both trimeric and pentameric HCMV complexes

After noting the significant increases in infection by TB40-BAC4_epi_ compared to TB40-BAC4_fib_, we hypothesized that cells within the organoids might express higher levels of receptors for the viral pentamer complex (Nrp2, THBD, and CD46) compared those for the trimer complex (PDGFRα and TGFβIII) (**Fig. 3A**). Others have demonstrated the importance of PDGFRα in organoid infections (21). Analysis of RNA levels from our recently published studies using bulk RNA-seq (11) show substantially higher levels of Nrp2 and CD46 compared to other receptors based on FPKM values, which supports the hypothesis (**Fig. 3B**). To analyze cell surface protein levels, we completed flow cytometry on uninfected NPCs and 45d organoids. Dissociated cells from either cultured NPCs or organoids were analyzed for expressions of TGFbRII (TC), PDGFRa (TC), Nrp2 (PC), CD46 (PC), and THBD (PC) using AF488, APC, or AF405 for which the gating is shown in F**igure. 3C**. For each sample, 10,000 events were recorded, and biological replicates were averaged, and we show the data as percentage of single or dual positive cells for the indicated receptors (**Fig. 3D,E**). We observed roughly 50% of cells in the NPC population expressing receptors for the virion trimer with PDGRFα being seen in 10% more cells than TGFβRIII (**Fig. D**). Receptors for the pentamer, Nrp2, CD46, and THBD, were detected in approximately 60% of the population. When evaluating cells expressing receptors for both the HCMV trimer and pentamer complex, we detected similar percentages for PDGFRα-Nrp2 or -CD46, yet reduced percentages for THBD-TGFβRIII receptors. Repeating the analysis using cells from dissociated organoids, we observed similar distributions of the receptors on cells, but at percentages lower than those for NPCs (**Fig. 3E**). These studies demonstrate that entry receptors for both the trimer and pentamer complexes are ubiquitously expressed.

**Figure 3.**
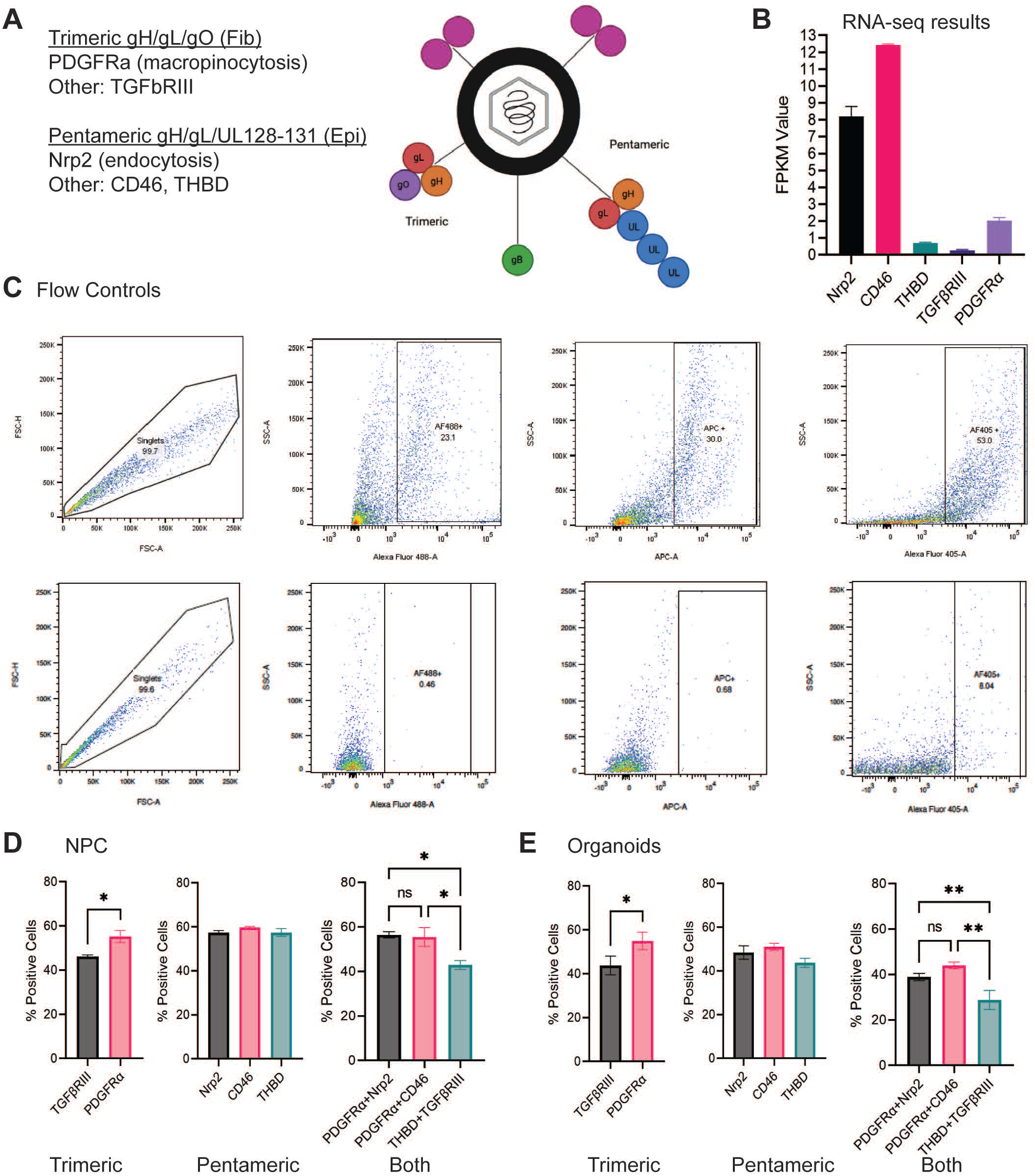
NPCs and cerebral organoids express several receptors required for HCMV entry. **(A)** Schematic depicting the HCMV viral particle noting the trimeric (gHgLgO) and pentameric (gHgLpUL128-131A) glycoprotein complexes and target cellular receptors PDGFRα and TGFβRIII (trimer) or NRP2, THBD and CD46 (pentamer). **(B)** RNA expression levels of these entry receptors as determined by bulk RNA-seq in cerebral organoids (11). **(C)** Gating strategy for flow cytometry assessing cell surface expression of entry receptors on iPSC-derived NPCs and cerebral organoids. Row 1 from left to right depicts traces from unlabeled cells, gate set for live cells labelled with AF488, APC, and AF405. Row 2 from left to right depicts traces from fluorescence compensation controls for the secondary antibodies. **(D)** Percentage of NPCs expressing the indicated receptors grouped by preferences or in combination. **(E)** Percentage of cells isolated from organoids expressing the indicated receptors grouped by preferences or in combination. Results were combined from 3 separate organoid differentiations and 4 NPC differentiations with at least 2 biological replicates per differentiation using one healthy control iPSC line for organoids and two for NPCs. Statistics were determined using students t-test for two groups or ANOVA with Tukey’s post hoc for comparison 3 or more groups.

HCMV has been shown to efficiently infect neural progenitor cells. Therefore, we evaluated NPC populations for expression of neural markers SOX2 and Tuj1 along with cellular entry receptors. To complement the cytometry studies, we analyzed expression using immunofluorescence. Uninfected, plated NPCs were stained using the indicated antibodies to entry receptors for trimer complex (**Fig 4A**) and pentamer complex (**Fig. 4B**) along with Tuj1 (β-tubulin III), Hoescht (DNA stain), and SOX2. In all samples, Tuj1 exhibits cytosolic staining consistent with microtubule localization, and SOX2 is colocalizing with the DNA stain. The entry receptors TGFBIII (**Fig. 4A**) and Nrp2, CD46 and THBD (**Fig. 4B**) exhibit staining on the periphery of cells likely indicating cell surface expression. Receptor PDGFR staining (**Fig. 4A**) is less clear with possible cytosolic and surface expression. The reduced number of cells staining for THBD is consistent with the flow cytometry analysis (**Fig. 3**). Finally, we determined the percentage of cells exhibiting SOX2 expression in NPCs and organoids using flow cytometry. A higher portion of cells within the NPC cultures, nearly 80%, expressed SOX2 compared to 50% in organoids (**Fig. 4C**), which is consistent with increased cell type diversity within the tissues. Co-expression of SOX2 with viral entry receptors ranged between 40-50% of NPCs (**Fig. 4D**) with cells expressing PDGFRα, Nrp2, and CD46 being in greater abundance. These percentages were reduced in cells isolated from organoids, which ranged from 30-40% of cells (Fig. 4D) reflecting the reduced SOX2 numbers. In summary, HCMV entry receptors for both the viral trimer and pentamer complex are expressed in both NPCs and cells from organoids, including SOX2 positive progenitor cells. Further, these data suggest that the increased tropism of TB40-BAC4epi compared to TB40-BAC4_fib_ is likely unrelated to receptor expressions and related to intrinsic features of the virion and associated entry process.

**Figure 4.**
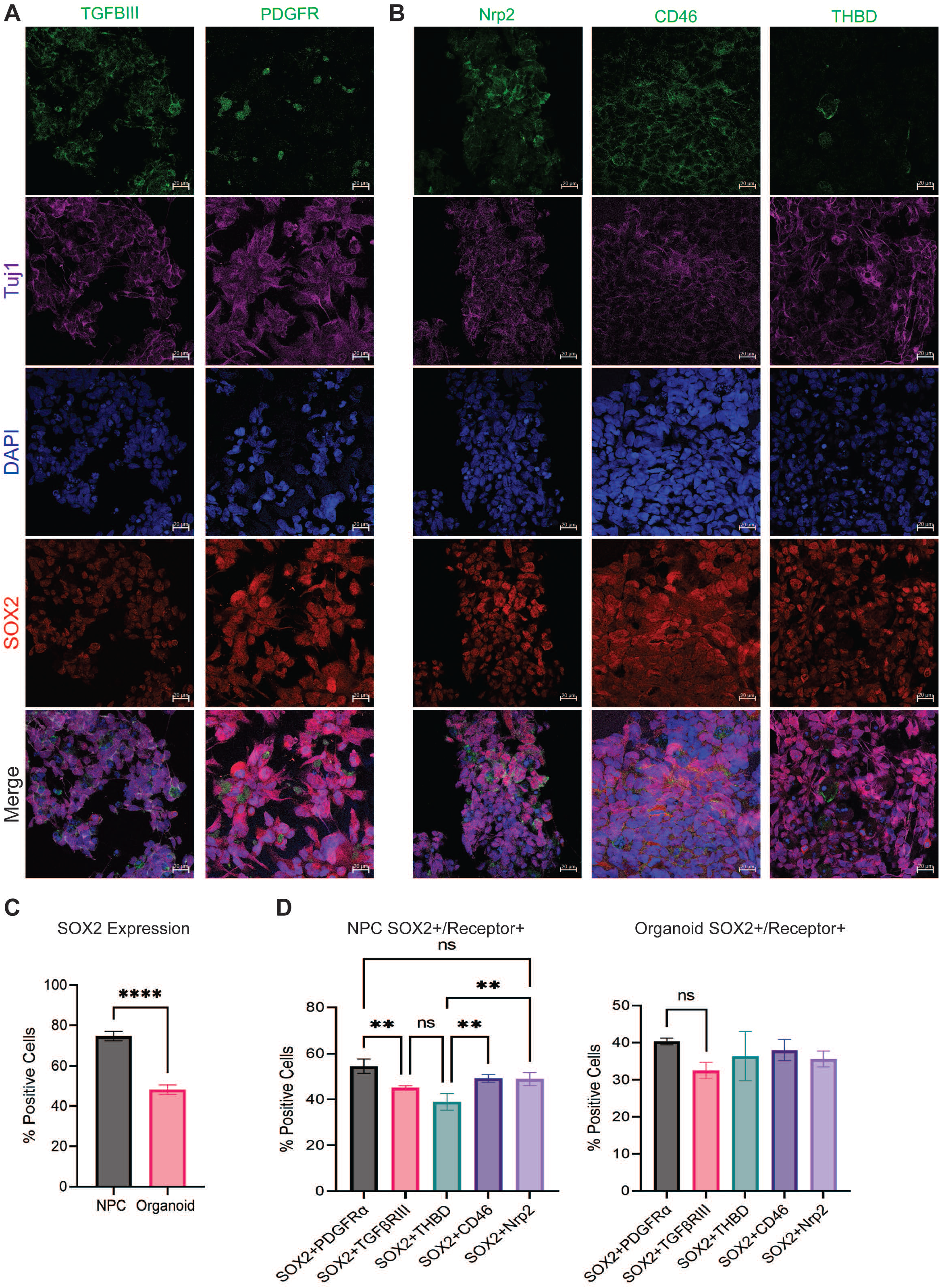
Viral entry receptors are expressed on SOX2-positive cells. Staining of staining panel of passage 3 NPCs 3 days post plate down labeling for **(A)** trimeric receptors and **(B)** pentameric receptors in green, neuronal beta-tubulin Tuj1 (purple), Hoescht (blue), progenitor marker SOX2 (red), and merged. **(C)** Percentage of cells expressing the progenitor marker SOX2 in NPCs and organoids as determined using flow cytometry. **(D)** Percentage of cells co-expressing SOX2 with entry receptor in NPCs and organoids. Statistics were determined using students t-test for two groups or ANOVA with Tukey’s post hoc for comparison 3 or more groups.

### Anti-HCMV neutralizing antibodies limit infection of cerebral organoids

Our end goal is to define HCMV neuropathogenesis and identify approaches to limit it using human neural tissue models. Considering that tropism is aligned with intrinsic features of the virions, we tested the impact of antibodies against viral glycoproteins previously shown to be neutralizing (15, 22, 23). We pretreated NPCs for 1 hr prior to infection with antibodies against HCMV gB or gH at 4 µg/ml. We also tested antibodies against host receptors PDGFR, previously shown to be neutralizing (21) and Nrp2. Subsequently, we infected with TB40-BAC4_epi_ at an MOI of 1 IU/cell in the presence of antibody for 2 hr after which virus and antibodies were removed and washed with PBS. Under these conditions, anti-gB and gH reduced cell-associated viral genome levels with the largest reduction occurring upon neutralizing gB (**Fig. 5A**). We did not detect reductions in viral genomes when using antibodies against cell surface receptors (data not shown). We evaluated the ability of anti-HCMV antibodies to limit infection per cell using immunofluorescence and staining the different conditions at 48 hpi. In untreated infections, we observed most cells expressing HCMV IE1 (**Fig. 5B**) including evidence of multinucleated syncytia with a central cytosolic assembly compartment. These infected cells exhibited enlarged nuclei, which is a hallmark of infection, and diffuse SOX2 staining. In contrast, neutralization using either anti-gB or anti-gH antibodies substantially limited the number of GFP and IE1 positive cells while maintaining nuclear SOX2 (**Fig. 5B**). Finally, we quantified changes in viral gene expression focusing on neutralization with anti-gB antibodies. We measured expression of the immediate early gene UL123 (i.e., IE1 protein), early gene UL44, and late gene UL99. The addition of anti-gB during infection of NPCs reduced expression levels for all gene classes at all time points (**Fig. 5C**).

**Figure 5.**
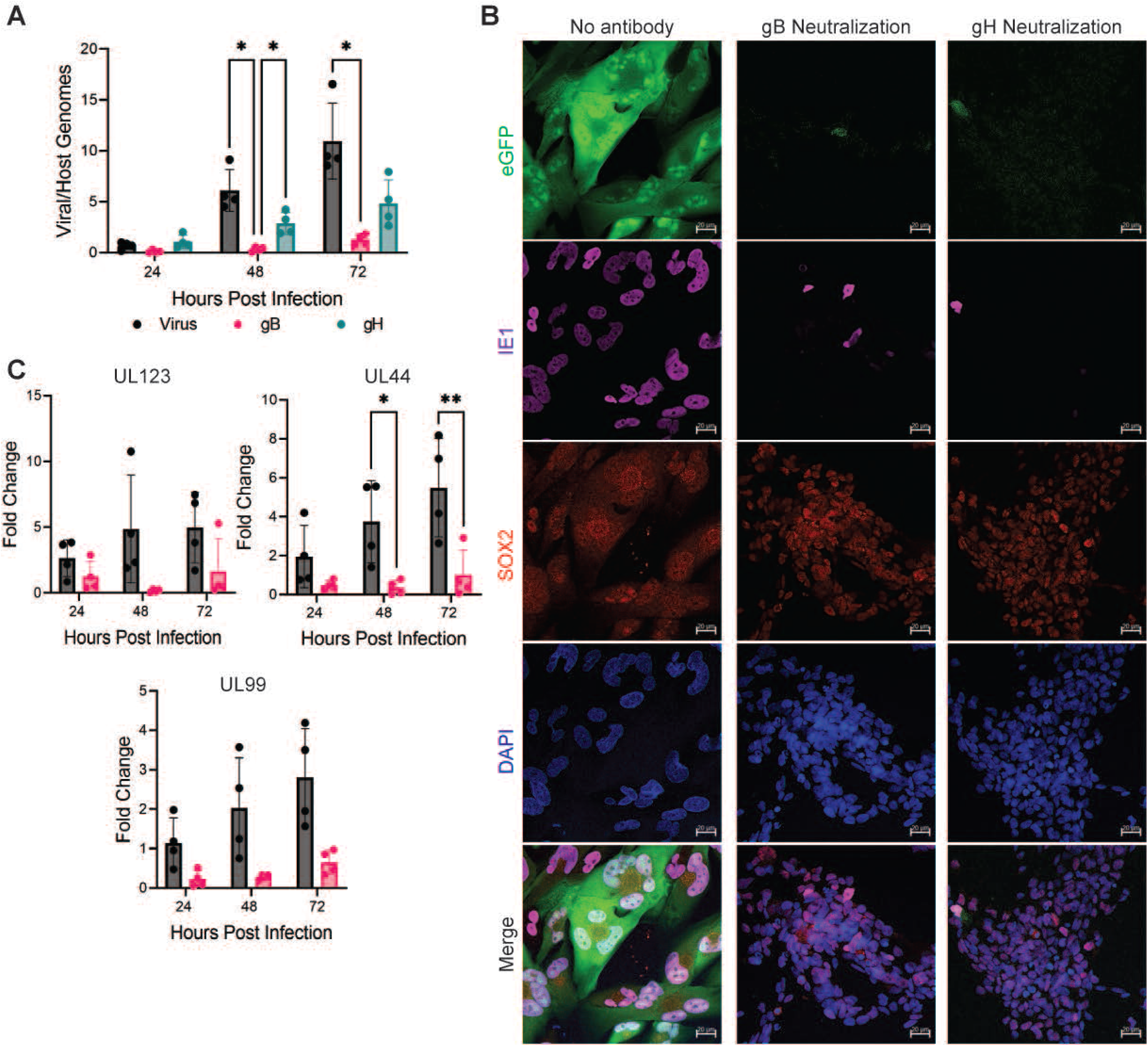
Anti-HCMV neutralizing antibodies limit infection of cerebral organoids. Plated NPCs were pretreated with the indicated antibodies then infected using TB40-BAC4_epi_ at an MOI of 1 IU/cell. After 2 hpi, cultures were washed with PBS and analyzed at the indicated times. **(A)** Levels of viral genomes relative to a cellular gene were determined using qPCR using whole cell DNA isolated at 24, 48, and 72 hpi (n=4). **(B)** Images of immunofluorescence staining done at 48 hpi on infected NPC or infected and with either anti-gB or anti-gH antibodies. Staining includes TB40-BAC4_epi_ eGFP (green), anti-IE1 (purple), SOX2 (red), and Hoechst (blue). All images were taken at 40x using the Zeiss LSM9800 microscope. **(C)** Quantification of viral transcripts was completed using RT-qPCR and primers to UL123, UL44, and UL99 relative to GAPDH (n=4). Statistical significance was determined using students T test.

Infection by HCMV causes IE1-dependent and -independent changes in neurodevelopmental gene expressions. We and others have demonstrated that IE1 mediates STAT3 nuclear localization contributing to deregulating SOX2 (24-27) To determine whether the anti-gB antibody can limit these pathogenic evens, we evaluated both changes in STAT3 localization and changes in expression of a subset of developmental genes. We measured changes in STAT3 localization by western blot analysis using nuclear fractions of NPC at varying times post infection. We controlled for fractionation by probing for nuclear Lamin B/C (**Fig. 6A**). We completed infections as described above by inoculating NPCs in the presence of TB40-BAC4_epi_ with and without anti-gB antibodies then analyzed total nuclear STAT3 along with phospho-STAT3 at multiple time points (**Fig. 6B**). We detected increased nuclear total STAT3 and reduced phospho-STAT3 during HCMV infection, which were near mock levels in the presence of neutralizing antibody. We quantified changes in total STAT3 and phospho-STAT3 normalized to total protein from multiple biological replicates (**Fig. 6C**). Next, we analyzed expression of neurodevelopmental gene targets FOXG1, FEZF2, DMRTA2, and EMX1. We have previously shown these to be downregulated within organoids following HCMV infection (11) and are key regulators in several developmental pathways involved in neurogenesis, cortical layer development, and progenitor maintenance (28-31). Infection of NPCs in the absence of gB antibody resulted in reductions compared to mock conditions in all four genes with significance by 24 or 48 hpi (**Fig. 6D**). In contrast, expression levels were similar to mock upon the addition of the neutralizing antibody. These data demonstrate that anti-HCMV neutralizing antibodies limit infection of NPC while preserving several changes known to be pathogenic in nature.

**Figure 6:**
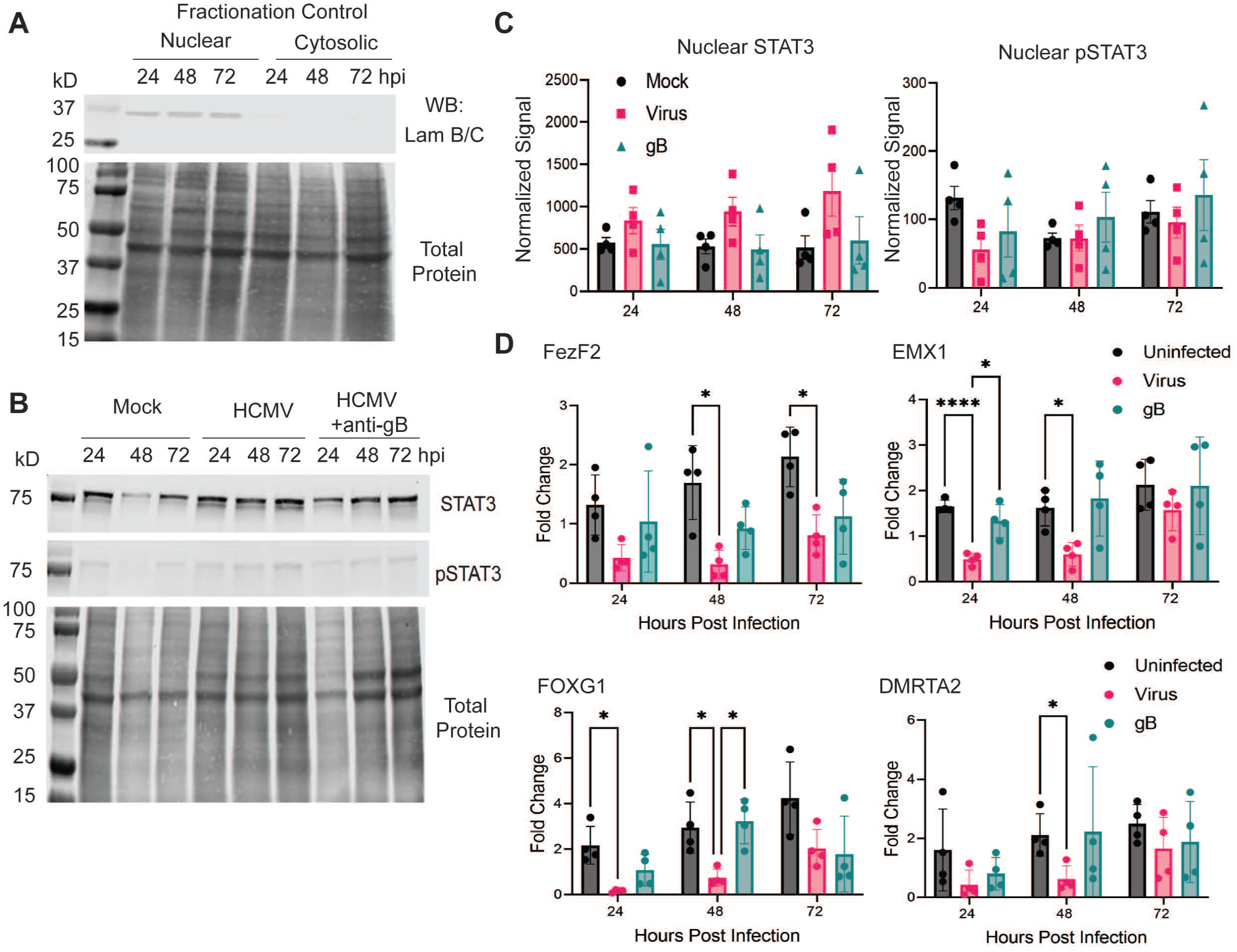
Neutralizing antibodies reduce HCMV-mediated disruption of developmental transcription factors. **(A)** Infected NPCs were isolated and fractionated into nuclear and cytosolic fractions. Fractions were analyzed by western blot analysis using an antibody to nuclear lamin A/C. Total protein is shown. **(B)** NPCs were mock-treated, infected using TB40-BAC4_Epi_ at an MOI of 1 or infected with the inclusion of an anti-gB antibody. Nuclear extracts were isolated and analyzed for total STAT3 and phospho-STAT3 at the indicated time points. **(C)** Quantification of western blots total nuclear STAT3 and pSTAT3 along normalized to total protein between wells from two iPSC lines across a total of four differentiations, two per line (n=4). **(D)** Analysis of FOXG1, FEZF2, EMX1, and DMRTA2 gene expression at 24, 48, and 72 hpi. Total RNA was collected, reverse transcribed and measured using primers to the specific genes. Samples were normalized to GAPDH, and the data are from two iPSC lines across a total of four differentiations, two per line (n=4). Statistical significance was determined using ANOVA with Tukey’s post hoc comparison.

In our previous studies, we found that maribivir treatment had only a modest effect on organoid structure and NPC function, so had proposed that the addition of a neurotrophic factor might offset some pathogenic consequences of infection (32). While the anti-gB antibody sustained FOXG1, EMX1, and FEZF2 expression immediately following infection in NPCs, the effects on Fez2 and FOXG1 were transient and lost by 72 hpi. Thus, we devised an alternative strategy combining the entry-blocking benefits of gB treatment with the neurotrophic factor brain derived neurotrophic factor (BDNF). BDNF has been shown to have an important role in cell survival, neurogenesis, and neuronal differentiation (33-37), and it is downregulated by murine CMV (38) and during HCMV-infection of cerebral organoids (11). We treated mock and HCMV-infected NPCs with either 4 µg/ml (high) or 1 µg/ml (low) concentration of anti-gB antibody in combination with BDNF at 20 ng/ml. Addition of neutralizing antibodies at both concentrations significantly reduced HCMV UL123 and UL44 RNA expression at all time points (**Fig. 7A**). A small drop in expression was observed in BDNF-only samples at 72 hpi. These differences were in line with changes seen by GFP fluorescence (**Fig. 7B**). Across the neurodevelopmental targets EMX1, FezF2, and FOXG1, we found that the combined treatment of anti-gB antibody and BDNF treatment maintained significantly higher target gene expression in HCMV-infected conditions at multiple time points post-infection compared to no treatment (**Fig. 7C**). In summary, addition of BDNF provided additional protection to anti-HCMV neutralizing antibodies in limiting infection-mediated disruption of several key neurodevelopmental genes in NPCs.

**Figure 7.**
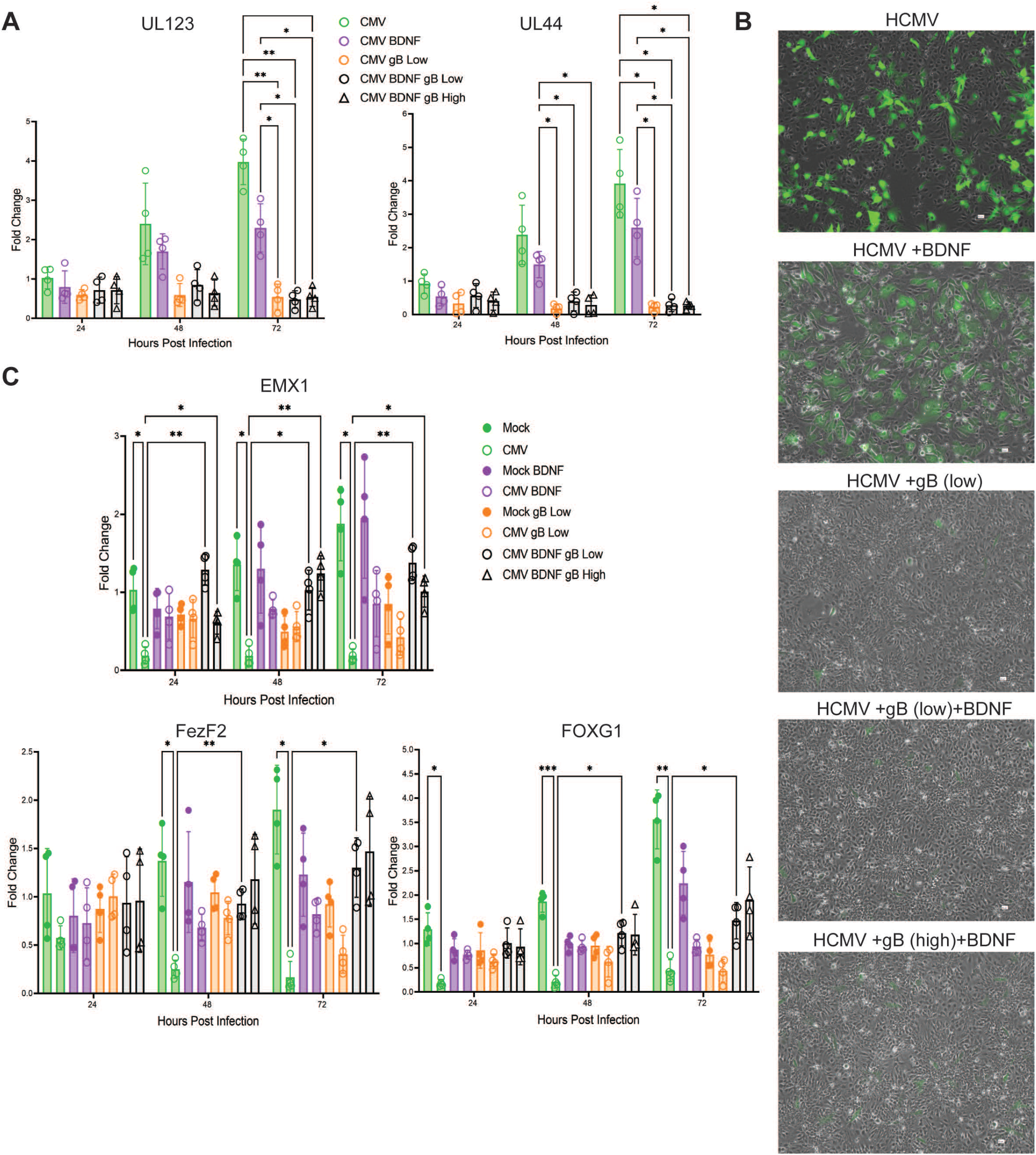
Neutralizing antibodies with BDNF provide significant protection against HCMV-mediated disruption of developmental factor expressions. NPCs were mock-treated, infected using TB40-BAC4_Epi_ at an MOI of 1 or infected with the inclusion of an anti-gB antibody at 4 ug/ml (high), 1 ug/ml (low) or BDNF at 2 ng/ml and analyzed at the indicated time points. **(A)** Quantification of viral transcripts was completed using RT-qPCR and primers to UL123, UL44, and UL99 relative to GAPDH (n=4). **(B)** Merged images of brightfield and GFP fluorescence of live cells taken 72 hpi from the various treatment conditions. **(C)** Analysis of FOXG1, FEZF2, and EMX1 gene expression at 24, 48, and 72 hpi. Total RNA was collected, reverse transcribed and measured using primers to the specific genes. Samples were normalized to GAPDH. These results were collected from two WT iPSC lines across a total of four differentiations, two per line (n=4). Statistical significance was determined using ANOVA with Tukey’s post hoc comparison.

## DISCUSSION

We and others have shown that HCMV infection induces significant disruption of neural development and function (4, 10-12). Previous data indicate that repeated passaging of viral stocks on fibroblasts results in increased accumulation of the trimeric versus pentameric components on the surface of the viral particle. In contrast, virus passaged on epithelial cells retain a more balanced composition (39). We found that infection using BAC-derived epithelial propagated virus TB40-BAC4_epi_ resulted in a significant increase in GFP fluorescence and viral DNA compared to infections using fibroblast propagated virus TB40-BAC4_fib_. In addition, the spread of fluorescence was significantly more in 3D tissues infected with TB40-BAC4_epi_ and likely includes syncytial formation as observed in infection of monolayer NPCs. We initially hypothesized that this difference in infection efficiency could be explained by an increase in the cellular receptors for the virion pentamer complex. However, contrary to this hypothesis, our analyses using flow cytometry and immunofluorescence determined that NPCs and cells from cortical organoids express similar levels of pentameric receptors (i.e., Nrp2, CD46, and THBD) compared to trimeric receptors (i.e., PDGFRα and TGFβRIII). Although at reduced percentages, similar disruptions were found on SOX2-positive progenitor populations. These data suggest that HCMV tropism for neural tissues is driven by the source of virus in contrast to the disruption of entry receptors.

Following virion attachment to the entry receptors, HCMV requires glycoprotein B (gB) to promote vesicle fusion with the host cell membrane (9, 13, 16). Previous studies have found that neutralizing antibody against viral glycoproteins can limit the spread of the virus and might be viable therapeutic targets (14, 40). Specifically, treatment with gB antibodies was shown to block *in vitro* infection of both epithelial cells and fibroblasts as well as blocking congenital infection in a guinea pig model (23), whereas experiments using antibodies to gH had mixed neutralizing activity based on the epitopes targeted by the antibody and the cell type of interest (14). With this in mind, we tested the capacity of gB and gH neutralizing antibodies to limit infection. Addition of antibodies at the time of infection significantly reduced viral DNA levels and gene and protein expression. Neutralizing antibodies partially protected NPCs allowing near mock levels of expression of neurodevelopmental transcription factors EMX1, FOXG1, DMRTA2, and FEZF2 supporting the importance of continued efforts in developing a therapeutic vaccine and neutralizing antibodies.

It is reasonable to suggest that complementary therapeutic approaches can be used to improve the health of the neural tissues during infection. In this regard, we focused on BDNF because of its known neurotropic properties including promoting neuronal differentiation, survival, metabolism, synaptic function, and repair. BDNF is currently being tested as an intervention to support neuron health in neurological diseases including Alzheimer’s disease (41, 42). Moreover, previous studies have shown significant downregulation of BDNF expression with CMV infection (11, 38), which could contribute to infection-induced neuronal damage (33). The inclusion of BDNF with anti-HCMV neutralizing factors showed additional protection of neurodevelopmental gene expression over antibodies alone. This dual treatment provides a potential therapeutic avenue for reducing viral entry as well as boosting neurodevelopmental transcripts.

In summary, we have demonstrated that epithelial-derived HCMV has increased tropism for neuronal tissues, and the addition of neutralizing antibodies in combination with neurotrophic factors provide protection in human neural cells and tissues.

## METHODS

### Cell culture and virus

Two independent control iPSC lines (4.2 and AICS-0057 (NPMI)) were grown and maintained in E8 as we have previously described (32). Line AICS-0057 was obtained from the Allen Institute for Cell Science. Following purchase, the line was thawed and expanded following our described procedure (32). Neural progenitor cells (NPCs) were cultured as a monolayer and differentiated using a Smadi Inhibition Kit from StemCell Technologies (#08581) requiring a minimum of 3 passages to become patterned NPCs. Cerebral organoids were generated using a cortical organoid kit from Stem Cell Technologies (#08570) that relies on an established protocol (19).

Viral stocks were generated using the HCMV TB40/E-BAC4 genome recombined to express the EGFP gene driven by an SV40 promoter (43, 44). Isolated BAC DNA was electroporated into MRC-5 fibroblast cells as previously reported (43, 44). The resulting virus was subsequently used to make high titered viral stocks by infecting MRC-5 fibroblasts resulting in virus TB40-BAC4_fib_ or by infecting ARPE-19 epithelial cells with viral stocks collected at passage 3 resulting in virus TB40-BAC4_epi_. For stock preparations, culture supernatants were cleared of cell debris, concentrated on a sorbitol cushion, and resuspended in media. For these studies, viral stock titers were determined on the cell line that they originated from, meaning TB40-BAC4_epi_ titered on ARPE-19 cells and TB40-BAC4_fib_ titers on MRC-5 cells. These concentrations were used in calculating MOI. Stock titers for TB40-BAC4_epi_ are similar between cells lines used for titering (MRC-5 2.3×10^8 IU/ml; ARPE19 1.8375×10^8 IU/ml; NPCs 2.425×10^8 IU/ml). In contrast, titers TB40-BAC4_fib_ is affected by the cell line used (MRC-5 5.35×10^8 IU/ml; ARPE19 5.65×10^7 IU/ml; NPCs 5.775×10^7 IU/ml).

Organoids on day 30 were infected with HCMV using 500 infectious units/ug of virus for 2 hours on a rocker and was generated from 3 separate differentiations using one WT iPSC line. Following infection, media was changed, and organoids were washed with PBS after infection media was replaced every 2 days. At 14 days post-infection, organoids were either fixed for cryosectioning or dissociated in Accutase enzyme and lysed for protein or DNA/RNA isolation. For NPC infections, P3 cultures and beyond were plated at 1 million cells per well and generated µfrom 4 differentiations using two different healthy control iPSC lines (4.2 and NPM1). The cells were then allowed to attach for 48 hrs prior to infection with HCMV TB40e-GFP virus at an MOI of 1. After a 2-hour infection period virus was removed, cells were washed with PBS, and fresh media was added. Following this, media was replaced daily. Cell pellets were then collected at 24, 48, and 72 hpi and processed for protein isolation or DNA/RNA isolation.

HCMV anti-gB human IgG1 (Absolute Antibody, Ab01475-10.0 Anti-HCMV gB [5A6]) and HCMV anti-gH human IgG1 (Absolute Antibody, Ab02066-10.0 Anti-HCMV gH [HCMV16]) were used for neutralization studies. Antibodies were added 1 hour prior to infection, followed by a wash with PBS; then fresh antibody was added during the 2-hour infection period. For studies using antibody and BDNF, the paradigm was the same except that BDNF (Peprotech, 450-02) was added along with the neutralizing antibodies for 1 hour prior to infection and for the 2-hour infection period. Following the infection, cells were pelleted and collected at 24, 48, and 72 hpi or fixed for staining. BDNF was used at a final concentration of 20 ng/mL; the antibody high concentration was 4 ug/mL, and antibody low was 1 u /mL.

### Immunofluorescence

NPCs were fixed on coverslips at various time points post-infection or without infection in 4% PFA and then permeabilized and stained. This was also the case for organoids which were fixed at 14 dpi or at 44 days of differentiation. Staining was then completed using these antibodies Nrp2 (R&D Systems, AF2215-SP), PDGFRa (BD Biosciences, 556001), CD46 (R&D Systems, AF2005), THBD (AbCam, ab33513), TGFβRIII (Sigma, T1940), UL123 (Shenk Lab, Clone 1B12), SOX2 (Sigma, AB5603MI), Tuj1 (GeneTex, GTX85469), and nuclear stain Hoescht (Thermoscientific, H3570). The appropriate fluorescent secondaries were then used in channels AF488, AF568, and AF647. Images were captured and produced using the Zeiss LSM980 (5x, 20x, 40x water) at various times post-infection.

### Flow Cytometry β

For organoids, the flow was run at day 45 of differentiation across 3 separate organoid differentiations with the same iPSC line and at least 2 biological replicates per differentiation. NPCs were analyzed after passage 3 and generated from 4 differentiations using the two healthy control iPSC lines. For each sample, 10,000 events were recorded, and biological replicates were averaged. Dissociation was done using Accutase (StemCell Technologies, 07920) enzyme to lift the monolayer of NPCs or dissociate the 3D structure of the organoid, the lysates were then filtered to remove debris and non-dissociated cells. After dissociation the cells were counted and prepped for flow using the same receptor antibody targets listed above (Nrp2, PDGFRa, CD46, THBD,

TGFβRIII) along with conjugated SOX2 (BD Biosciences, 561610). Secondaries of AF488 and AF568 were used in correspondence with the correct primaries. Samples were run on the LSRII machine with 10,000 events recorded per group. The graphics and subsequent analysis were conducted using the program Flowjo.

### Protein and nucleic acid analyses

DNA samples were collected at time points indicated post-infection from NPCs or organoids using Qiagen Kit (Qiagen, 69506). Following isolation, DNA PCRs were run using primer sets for UL123 and GAPDH then the ratio of Cq values was calculated and reported in the graphs as Viral/Host genomes. Total RNA was isolated (Qiagen, 74136), and reverse transcribed into cDNA using the Promega RT Kit (Promega, A3500). RT-qPCR was performed using specific primer sequences for FOXG1 (F-CGTTCAGCTACAACGCGCTCA, R-CAGATTGTGGCGGATGGAGTT), EMX1 (F-AGCCCCGTCTTAATGCAACA, R-CTAGGATTGCGGGGCTAGTG), FEZF2 (F- GTGGTGGAATTCGCCGCCGCCATGGCCAGCTCAGCTTCCCTGGAGACCATGGTG, R- TGCTGGATATCAGCTCTGAACTGTCCTGGCTAGGTCCTTTGCTGA), DMRTA2 (F- GCCTGCCTACGAAGTCTTTGGCTCGGTTT, R- CGTCTTGGGAAACAGATCAAACTTCTG), UL122 (F-ACCTGCCCTTCACGATTCC, R- ATGGTTTTGCAGGCTTTGATG), UL123 (F-GCCTTCCCTAAGACCACCAAT, R- ATTTTCTGGGCATAAGCCATAATC), UL44 (F-GCCCGATTTCAATATGGAGGTCAG, R- CGGCCGAATTCTCGCTTTC), and UL99 (F-GTGTCCCATTCCCGACTCG, R- TTCACAACGTCCACCCACC). The resulting Cq values were normalized to GAPDH (F- GTGGACCTGACCTGCCGTCT, R-GGAGGAGTGGGTGTCGCTGT).

Protein samples were isolated from cell pellets at 24, 48, and 72 hpi using a nuclear and cytoplasmic fractionation kit (Thermofisher Scientific (Cat# 78833)) and protease and phosphatase inhibitors (Thermofisher (Cat# 78425 & 78428)). Once the protein was isolated, the lysates were quantified using a BCA assay from (Thermofisher Scientific (Cat# A53225)). Western blots were run at 105 volts for 1 hour with 25 µg of protein sample loaded in each lane. To quantify total protein, the Revert total protein stain system was used (Revert Kit (LiCor, 926-11015)). After staining for total protein, blots were incubated in blocking buffer for 1 hour and probed using primaries Lamin B/C (Cell Signaling, 4777S), STAT3 (Cell Signaling, 9139S), and pSTAT3 (Cell Signaling, 9138S). After overnight incubation in primary, blots were incubated in appropriate secondary for 30 minutes using rabbit green Secondary (LiCor, 926-32213) and mouse red Secondary (LiCor, 926-68072). Finally, blots were imaged on LiCor LCX imager and quantified using image studio lite version 5.2.

### Statistical analysis

Conclusions were made based on data collected from three independent organoid differentiation and infections followed by one-way ANOVA or Student’s t-test as appropriate with Tukey post hoc test for DNA PCR, qPCR, and western blot. Data are presented as mean with SD and p<0.05 was considered significant.

## Acknowledgments

We thank Tom Shenk for providing antibodies against HCMV IE1, IE2, and pp28 proteins; Benedetta Bonacci in the Versiti-Blood Research Institute Flow Cytometry Core; and MCW Cancer Center and Children’s Research Institute’s Flow Cytometry Core. We thank Melissa Whyte and Halli Miller for their helpful advice on flow cytometry and members of the Terhune and Ebert laboratories for their input on the project.

Research reported in this publication was supported by the National Institute of Allergy and Infectious Diseases division of the National Institutes of Health under award number R01AI132414 (S.S.T. and A.D.E.). The content is solely the responsibility of the authors and does not necessarily represent the official views of the National Institutes of Health. These studies have also been supported by a generous philanthropic gift by The Stead Family Foundation to define the impact of infection and inflammation on brain health.

## Notes

### Competing Interest Statement

The authors have declared no competing interest.

